# Evolution uncovers a general tradeoff between recovery after heat shock and growth at elevated temperatures

**DOI:** 10.1101/2025.10.22.683249

**Authors:** Akshat Mall, Katelyn J. Rode, Christopher J. Marx

## Abstract

The maintenance of microbial diversity depends upon fitness tradeoffs between environments. While the importance of tradeoffs is clear, it has been surprisingly difficult to predict which traits they will occur between and at how granular a level. For example, it is unclear whether performance between a constant versus a pulsed exposure of the same stress tend to be positively correlated, independent of each other, or negatively correlated. Empirically, it has been shown that a critical feature structuring environments is temperature. However, the compatibility between strategies to deal with different forms of heat stress is unclear. For instance, are strains that grow well at higher temperatures also stronger at withstanding heat shock? To understand how environmental microbes can adapt to better deal with different forms of heat stress, we performed an evolution experiment using a dominant phyllosphere microbe *Methylobacterium extorquens* in a regime of intermittent heat shock. We identified the genetic basis of adaptation, discovering a large number of loci capable of mediating adaptation to heat shock, most of which previously had not been linked to heat stress. Despite the genetic divergence, we discovered a general tradeoff between heat shock resistance and growth at consistently elevated temperatures. We found this tradeoff was not limited to evolved isolates, but was also represented across a sample of environmentally isolated *Methylobacterium* strains. These findings indicate a generic conflict between strategies to deal with heat shock recovery and growth at elevated temperatures, suggesting even variation in intensities of a stressor can drive diversity in microbial strategies.

## Introduction

Microbial populations incessantly encounter a number of environmental stressors. Biochemical and resource constraints limit the ability of cells to mount responses to deal with all variations in the environment. These constraints can manifest as tradeoffs associated with different responses and drive diversification of microbial strategies, as is commonly observed in ecological communities^1,2^. While a number of studies have explored adaptive strategies and tradeoffs between distinct environmental stressors^3–6^, less is known about the challenges imposed by the same stressor when present continuously versus in a fluctuating manner, and how populations tackle these different challenges. One common stressor which presents itself in these distinct manners is high temperature which can be encountered at varying intensities and durations.

Temperature variation is one of the fundamental drivers of natural microbial communities^7^, shaping metabolic strategies^8,9^, community composition and function^10–12^, and adaptation^13,14^. Most studies on understanding how temperature structures microbial physiology and diversity have focused on a constant or average temperature. However, temperature can fluctuate sharply, not just seasonally, but even within a day. While it is evident different temperatures favor different strategies and can drive diversification^15,16^, it is unclear if or how the degree of variations in temperature shapes thermal traits. For instance, optimal strategies for thriving in “high temperatures” might differ for increased temperatures of different intensities and durations, and the direction or degree of correlation between these different strategies is not known. While both the magnitude and frequency of heat stress can be variable (Fig 1A), we focus on two extreme cases in this work – a constant but habitable increase in temperature (Fig 1B) and a short-lived but lethal increase in temperature followed by recovery at an optimal temperature (heat shock, Fig 1C). Understanding the factors shaping thermal traits of microbes is of critical importance in predicting adaptation and community dynamics in nature.

**Fig 1.**
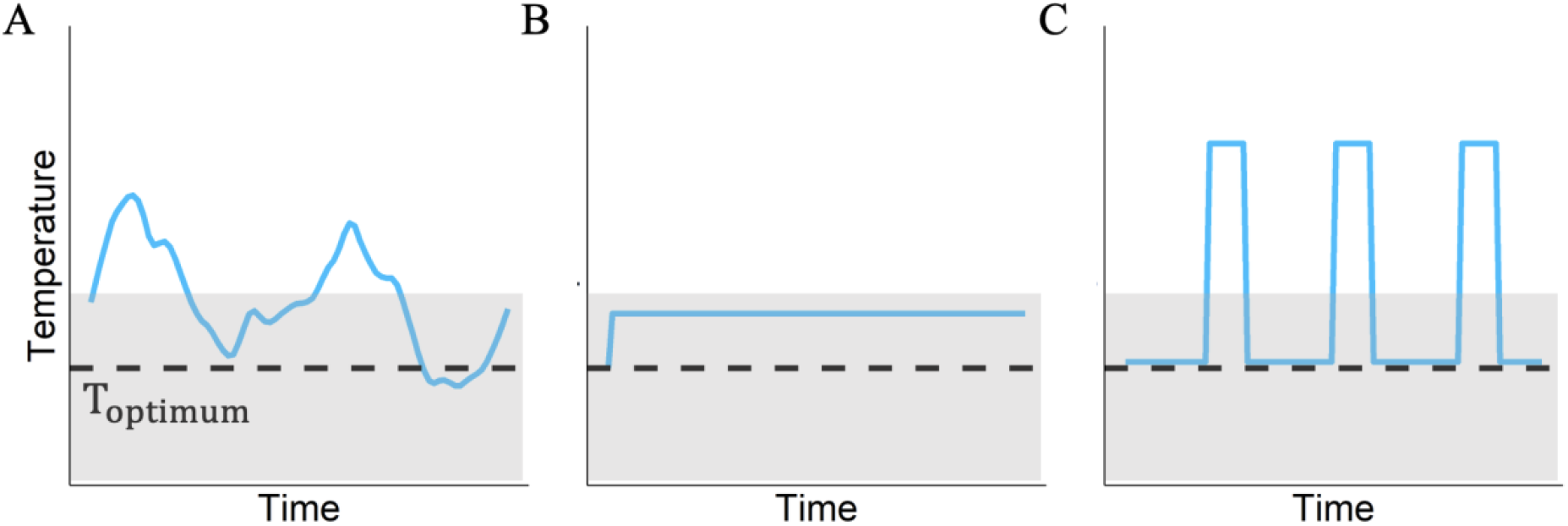
Possible variations of heat stress. Each panel represents a profile of temperature vs time (blue curve). Dashed black line represents the optimum growth temperature of the microbe being studied, and temperatures above the dashed line constitute heat stress. Gray shaded area represents habitable temperature range permitting growth while temperatures outside this zone are lethal. **A)** Pattern of heat stress observed in natural environments. **B)** Most studies on heat stress focus on a constant elevated temperature, as shown here by the horizontal blue line. **C)** Heat shock – the pattern of heat stress forming the primary focus of this work.

How microbes in natural ecosystems are affected by, and adapt to heat shock (Fig 1C) remains underexplored. Research on adaptation to increasing temperatures has focused largely on continuously elevated temperatures which enable growth^17–19^. The relatively few studies that focus on how bacterial populations can adapt to heat shock have been confined to pathogenic microbes, and our understanding of pleiotropic effects of improved heat shock resistance limited to phenotypes related to virulence^20–22^. Heat stress can be encountered by microbes in a number of contexts, such as stochastic fluctuations in the natural environment, forest fires, anthropogenic settings, etc. It is unknown how microbes in other environmental contexts deal with recovery from, and can adapt to, heat shock. Of particular interest is the potential synergy or antagonism between strategies to deal with heat shock and constantly elevated temperatures.

How strategies to deal with thermal stress of different durations and intensities affect each other is not clear. A primary detrimental effect of both continuously elevated temperatures and heat shock is increased generation of reactive oxygen species (ROS)^23,24^ and ROS removal helps alleviate the damaging effects in both instances^24–26^. Protein misfolding is another common impairment during thermal stress^27^, and the physiological response to different types of heat exposure involves upregulation of heat shock proteins and chaperones, and global stress response^28,29^. Overexpression of heat shock proteins is a mechanism facilitating adaptation to both growth at elevated temperatures and heat shock resistance^18,22,30^. Owing to these similarities, we might expect strains adapted to growth at high temperature to be better at surviving heat shock, and vice versa.

Alternately, a constant increased temperature that permits steady-state growth at a slower rate is distinct from surviving and recovering from a short duration of lethal heat stress, such that the optimal strategies to deal with either could differ. For instance, loss of function mutations in a key molecular chaperone *dnaJ* have been shown to improve heat shock resistance at the cost of growth at elevated temperatures in *Salmonella enterica* and *Escherichia coli*^21^. Whether such a tradeoff is idiosyncratic to *dnaJ*, and/or the microbes involved or hints at a broader conflict between strategies for dealing with heat stress of varying intensities is unclear.

Evolution experiments offer a tractable approach to understand the feasibility and diversity of routes enabling adaptation to an environmental shift. A number of studies across many organisms have explored how microbes can adapt to a consistently increased temperature, improving our understanding of the genetic targets and molecular mechanisms facilitating adaptation^19,31^ and of the patterns of pleiotropic effects associated with adaptation to growth at higher temperatures^32,33^. Notably, few mutations, sometimes just one^31^, are sufficient to provide significant fitness benefits at higher temperatures and allow for an expansion of the thermal niche. The pleiotropic effects associated with adaptation to increased temperature have largely been studied with respect to changes in thermal niche for growth, but not different intensities of thermal stress. While the average effect of adaptation to high temperatures is a fitness defect at lower temperatures, the observed tradeoffs have been idiosyncratic, with no significant correlation between the degree of fitness improvement in one environment with fitness defect in another^32^. We lack a similar understanding of the diversity of loci and mechanisms which allow adaptation to heat shock, and whether the mechanisms driving improved heat shock resistance aid or inhibit growth at elevated temperatures.

In this study, we use *Methylobacterium extorquens* PA1 to study adaptation to heat shock. *Methylobacterium* species constitute dominant members of the leaf microbiome^34^ and play critical ecological roles, from affecting plant health to influencing biogeochemical cycles^35–37^, and constitute model systems for a number of biotechnology applications^38–40^. Temperature variation is a major abiotic factor shaping microbial communities associated with plants^41,42^. *Methylobacterium* communities isolated from leaf surfaces display distinct ranges of optimum growth temperature, suggesting temperature as a critical factor shaping community diversity and composition^43^. This implication that temperature drives distinct growth strategies and thus diversity is similar to what is known for other microbes. However, like in other systems, it is not clear what role of temperature variation, and not just average temperature, drives distinct strategies.

We discovered that a major impact of heat shock on bacterial cells is not just loss of viability, but a significantly increased lag time amongst the survivors. Using evolution experiments with fluctuating exposures to heat shock, we found divergent adaptive outcomes which improve fitness by reducing the recovery time after heat shock. We found that the evolved isolates, despite distinct genetic bases, exhibited a strong-tradeoff between improvement in recovery after heat shock and growth at elevated temperatures. We discovered that the tradeoff was not limited to the evolved isolates but was also represented within a set of environmentally isolated *Methylobacterium* strains, suggesting an incompatible physiological distinction between strategies suited for heat shock resistance and growth at continuously elevated temperatures.

## Materials and Methods

### Bacterial strains and culturing

The ancestor strain used in this study is derived from *Methylobacterium extorquens* PA1^44^, and differs from the reference strain in a) mutations present in our laboratory stock and b) deletion of cellulose synthesis genes to avoid biofilm formation and clumping, which aids growth measurements in liquid culture. This *Δcel Methylobacterium extorquens* PA1^45^ is the WT (and ancestor) strain for this work. All cultures were grown in *Methylobacterium* PIPES (MPIPES) minimal media^46^ in volumes of 5 ml. Cultures were supplemented with either 15 mM methanol, or 3.5 mM succinate as a carbon source. For evolution experiments, and all experiments using evolved isolates, methanol was used as the substrate. For quantifying tradeoffs among environmentally isolated strains, both methanol and succinate were used as the substrate to probe the role of substrate in the observed tradeoff. For plating assays, agar plates were made with addition of 1.5% (w/v) agarose in MPIPES minimal media, and supplemented with 15 mM succinate as a carbon source. Growth conditions were 30 °C (unless otherwise stated) and shaking at 250 rpm. The environmental isolates used in Fig 5C are listed in Table S1.

### Evolution experiment

Ten individual WT colonies were randomly chosen and served as ancestors for each replicate population. These colonies were inoculated in 5 ml cultures (supplemented with 15 mM methanol) and allowed to grow for 3 days. After this period, 500 µl was taken from each replicate into an epitube and placed in a 55 °C water bath for 5 min. After heat shock, 78 µl was transferred into fresh 5 ml cultures, and this serial transfer regime was repeated 20 times. Every 3 transfers, subpopulations of each replicate was frozen at -80 °C, with 8% DMSO as a cryoprotectant.

#### Viability analysis

WT cells were grown in 5 ml cultures with 15 mM methanol until stationary phase. From this culture, 500 µl was placed in an epitubes and heat-shocked by placing in a water bath at 55 °C for different durations of time (Fig 2A). To assay viability, cells were spot-plated onto MPIPES agar plates (with 15 mM succinate) at different dilutions, following the protocol in Lee et al^47^.

**Fig 2.**
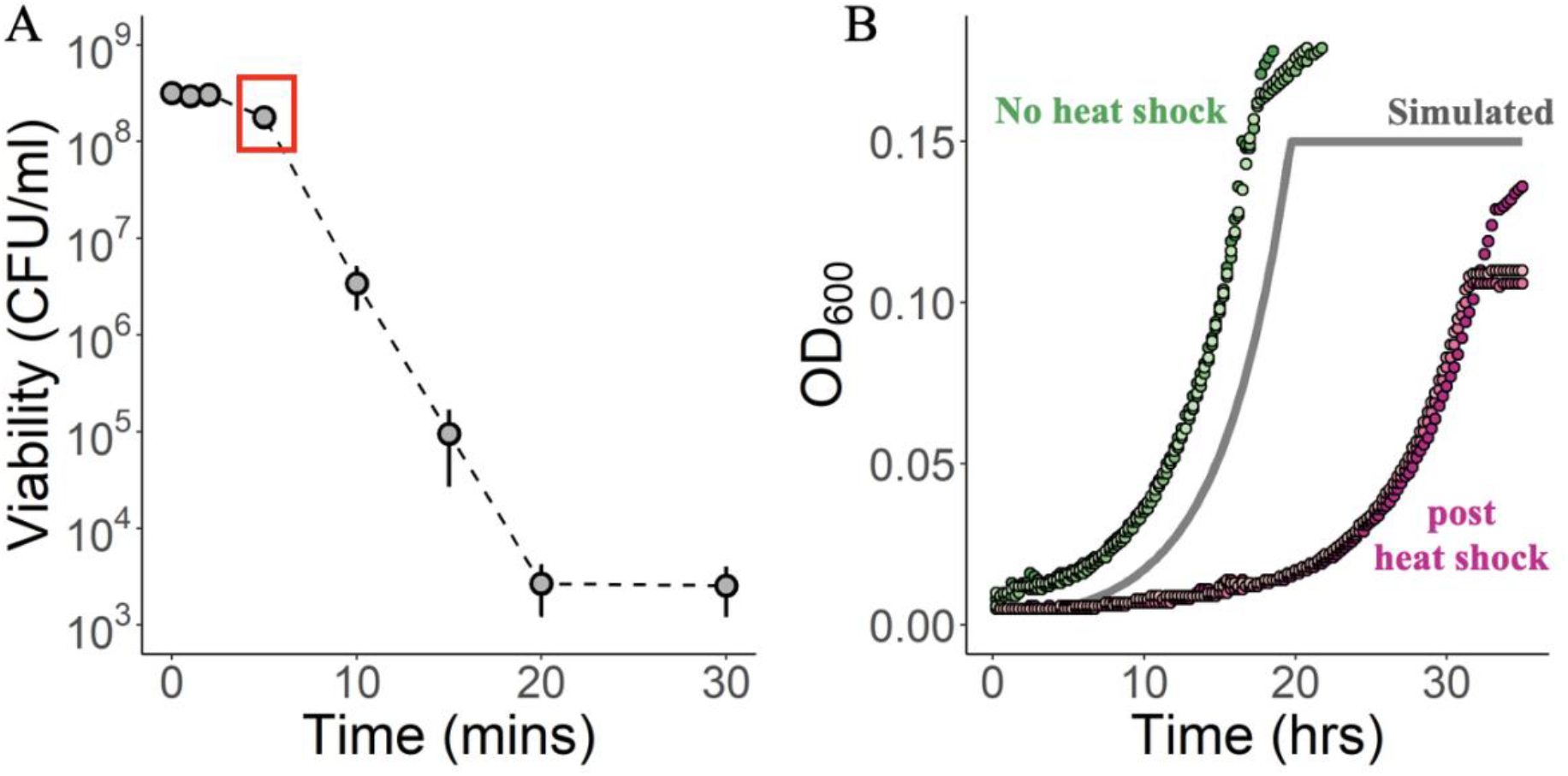
Heat shock leads to A) loss of viability and B) increased lag time. **A)** Number of viable cells vs duration of heat shock at 55 °C. Each point represents the mean of 3 biological replicates, and error bars denote standard error of mean. Error bars are only visible when bigger than plotting symbols. Point enclosed in red square denotes the regime (5 min) used for further analysis in panel B, and later evolution experiments. **B)** Growth curves shown on a log_10_ scale (3 replicates each) for populations after (pink) or without (green) heat shock at 55 °C for 5 min. Gray (simulated) curve represents the expected growth curve if the only impact of heat stress was a loss of viability as estimated from panel A (point enclosed by red square).

### Growth analysis

Growth data was quantified by growing cultures in a 48-well plate (Corning) in a Biotek Synergy H1 Plate Reader (Agilent Technologies). When grown in multi-well plates, shaking speed was set at 548 rpm in a double orbital motion, and culture volume was 640 µl. Growth curves were created from the raw data with a custom R script (see Data Availability). Growth rates and lag times were quantified using Qurve^48^. In brief, a line was fit to log-transformed OD vs time data, and the maximum slope over a 3h or longer period (at least 6h from the start of the experiment) was designated maximum growth rate. This line was extrapolated and intersection with the starting OD was used to quantify lag time (see Fig S1 for an example).

### Genome sequencing and analysis

After 20 transfers, and ∼140 generations of evolution, colonies from each replicate were plated out on individual agar plates supplemented with 15 mM succinate. From each replicate, 1 or 2 colonies were chosen for whole genome sequencing. In the instances where 2 colonies were chosen, it was because there was visible heterogeneity in colony size or morphology on that plate. The chosen colonies were grown in a 5 ml culture (with 15 mM methanol) until stationary phase. DNA was extracted using MasterPure Complete DNA and RNA Purification Kit (Biosearch Technologies) following the manufacturer’s protocol for bacterial samples. For all isolates, whole genome sequencing was conducted on the Illumina NextSeq2000 platform by Seqcoast Genomics (Portsmouth, NH, USA). For isolate A1, genome sequencing was also conducted using Oxford Nanopore Technology by Plasmidsaurus. For each isolate, mutations were identified using breseq^49^ by comparing reads against the ancestral genome. The WT ancestor used in this work differs from the NCBI GenBank reference genome (accession no. NC_010172) for *Methylobacterium extorquens* PA1 by 6 mutations (Table S2).

## Results

### Heat shock leads to an increase in lag time

We hypothesized that, besides cell death, heat shock could result in damage and an increased recovery time for even the cells which survive^50^. In order to understand the impact of heat shock on bacterial cells, we first characterized the kill curve for heat shock at 55 °C. We observed a decline in number of viable cells with increasing durations of exposure to heat stress (Fig 2A), as seen previously for other microbes^23^. We next focused on the regime of a short exposure to heat (5 min at 55 °C) where populations suffered relatively little loss of viability, and observed that populations experiencing transient heat stress experienced a long lag, much longer than could be explained by cell death (Fig 2B). Populations diluted into fresh media without heat treatment (green) exhibited little, if any, lag. However, populations diluted into fresh media after 5 min of heat stress at 55 °C (purple) exhibited a long lag. This extra lag is not just a consequence of cell death, where the number of growing cells is reduced and surviving cells go through extra generations to reach the same optical density. Estimating the number of survivors after a transient heat shock, from Fig 2A, we simulated a growth curve (gray) accounting for cell death, and found that an approximately two-fold loss of viability could not explain a major portion of the extra lag. The initial impact of a short duration of heat stress to *M. extorquens* is fairly minor upon cell viability, but generates a sizable increase in lag time. This suggests that cells which are able to recover after heat shock do so with a longer lag, due to either damage or a physiological response to the heat stress.

### Evolution to cycles of heat shock selected for diverse beneficial mutations

To study how *Methylobacterium extorquens* can adapt to improve fitness in an environment punctuated with heat shock (Fig 1C), we evolved 10 replicate populations of wild-type *Methylobacterium extorquens* PA1 on a cyclical environment of growth on methanol at 30 °C and a transient heat shock at 55 °C for 5 min, followed by dilution and re regrowth on methanol (Fig S2). After 20 cycles or ∼140 generations, we performed whole genome sequencing for 14 isolates (either 1 or 2 isolates per replicate population). Despite some parallelism, we found many different genetic targets of adaptation (Fig 3, Table S3 for mutation details). This is in contrast to previous studies of evolution on heat shock, which found evidence of highly convergent adaptation where all isolates were observed to have mutations in the same locus^21,22^. Of the observed genetic targets of adaptation, three loci had mutations in more than one isolate. These loci were *secA, dnaJ*, and *hsp20* (for *hsp20* – same family of proteins but different gene each time). Peculiarly, despite sequence analysis via breseq^49^ targeting SNPs, indels, new junctions, and copy number variations, we failed to find any mutations in isolate A1.

**Fig 3.**
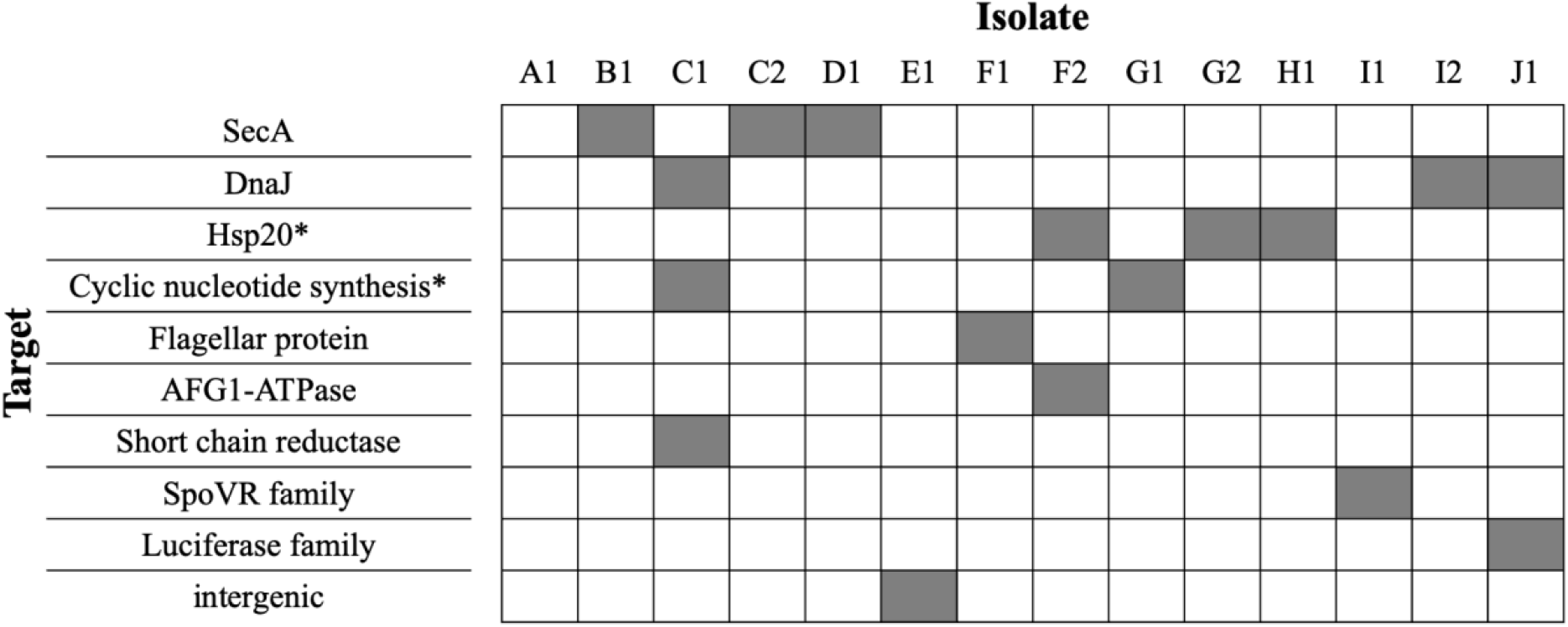
Mutations identified in evolved isolates. Isolate name denotes replicate information (A-J), and the number (1 or 2) designated to the isolate picked for that replicate. E.g. – C1 and C2 denote different individuals from the same replicate in the evolution experiment. Locus name denotes the class of genes or functions the mutation was predicted to affect. Boxes are filled when an isolate had a mutation in that genetic loci. Asterisk (*) near locus name indicates that the mutations in the different isolates occurred in different homologs within the same gene class.

The diversity of mutational targets observed in our evolution experiments hint at a large number of different accessible routes for adaptation to heat shock. Of the observed targets of adaptation, two loci, *dnaJ* and *hsp20*, are directly known to play a role in dealing with heat stress. However, the roles of other genetic targets in helping deal with heat stress remain unclear.

### Evolved isolates exhibit reduced lag after heat shock

The lag time and growth rate following heat shock were assayed for the evolved isolates in order to determine which of these fitness components exhibited an improvement. We characterized the performance of the evolved isolates in the selection regime of heat shock at 55 °C for 5 min followed by growth on methanol at 30 °C. Only one evolved isolate had a significant change in growth rate after heat shock (Fig 4A), and it was slower. On the other hand, all but two isolates exhibited a significant decrease in lag time after heat shock (Fig 4B, Fig S3).

**Fig 4.**
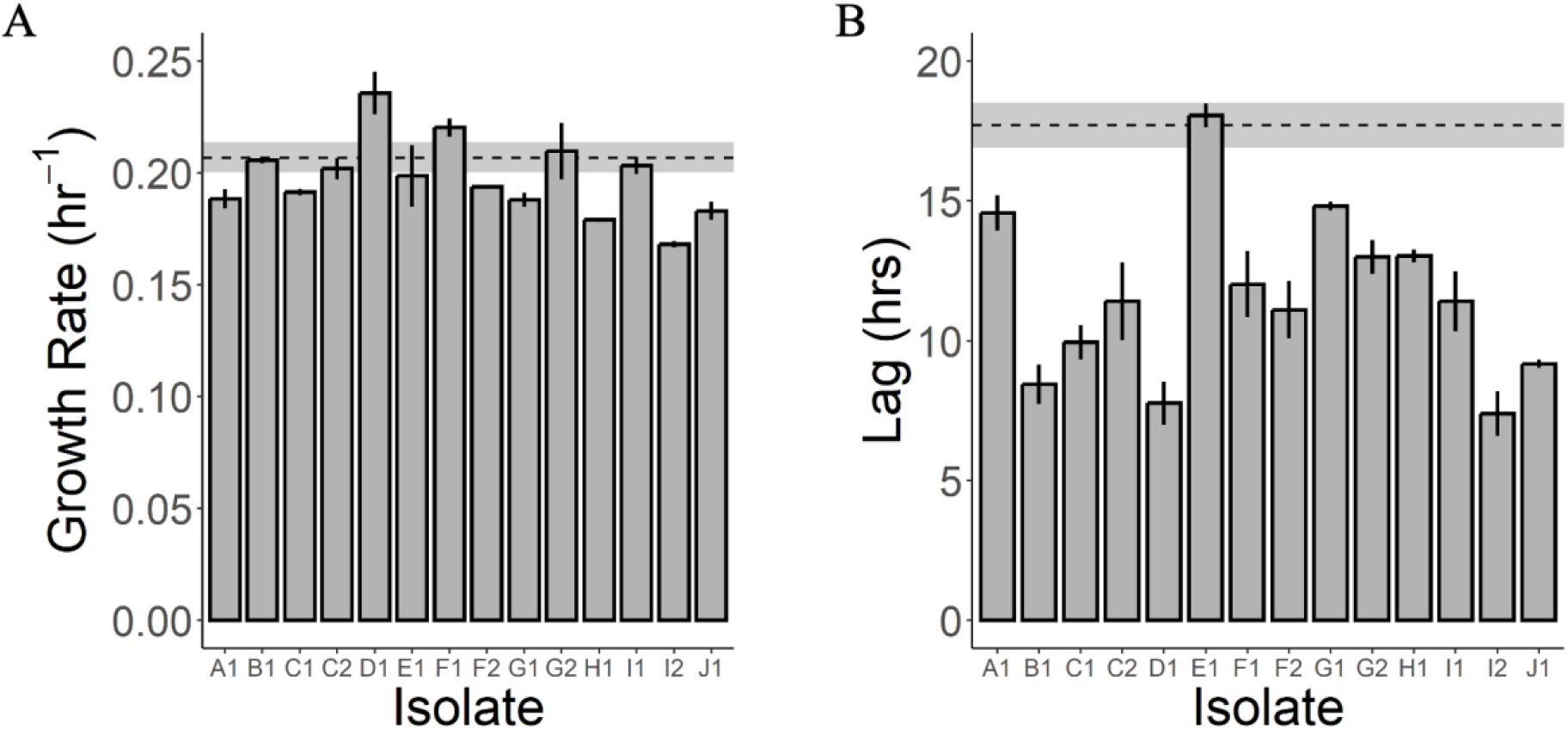
Evolved isolates A) do not exhibit any consistent change in maximum growth rate but B) do exhibit reduced lag time after heat shock. In both panels, bars show the mean of 3 replicates, and error bars denote standard error of mean. Dashed line and shaded region represents mean and mean ± standard error for ancestor. **A)** The change in maximum growth rate is not statistically significant (*p* > 0.05, Student’s t-test) except for I2 (*p*=0.024). **B)** The decrease in lag time is statistically significant (*p* < 0.05, Student’s t-test) for all isolates except E1 (*p*=0.69) and G1 (*p*=0.065).

**Fig 5.**
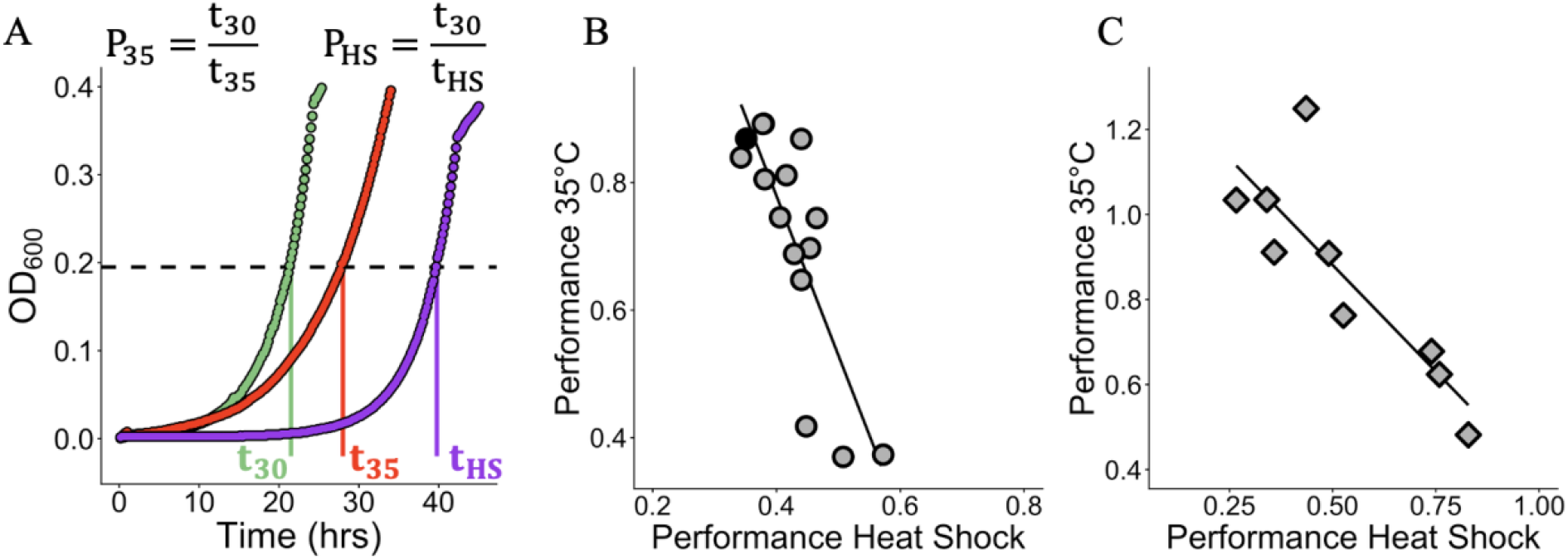
General tradeoff between heat shock resistance and growth at elevated temperatures. **A)** To quantify the tradeoff between growth at 35 °C and recovery after heat shock, we define a metric “performance” (P_x_) which is the time taken to reach a threshold optical density (black dashed line) during growth in a normal regime of 30 °C relative to the regime of interest. A performance of 1 indicates time taken to reach threshold OD was the same as the time taken at 30 °C. A performance of 0.5 indicates the isolate took twice as long to reach the threshold OD relative to time taken at 30 °C. We use this metric instead of other common metrics like growth rate because many isolates fail to follow an exponential or logistic growth dynamic in high temperature regimes (see Fig S2 for examples). **B)** Performance during growth at 35 °C vs performance in a heat shock regime for evolved isolates (gray circles) and ancestor (black circle). Each point represents the mean of 3 replicates. Black line represents the line of best fit (R^2^ = 0.66). **C)** Performance during growth at 35 °C vs performance in a heat shock regime for 9 environmentally isolated *Methylobacterium* strains. Each diamond represents the mean of 4 replicates. Black line represents the line of best fit (R^2^ = 0.72).

We discovered two isolates with surprising effects. First, isolate A1 had no detectable mutations, and yet was more fit than the ancestor, exhibiting a significantly shorter lag after heat shock. Since the variant calling was a hybrid of short and long reads, this rules out rearrangements or copy number variation. As such, our current hypothesis is that this represents an epigenetic adaptation. Further work will be required to determine if this is caused by alternative DNA methylation or some other mechanisms^51,52^. Second, isolate E1 harboured a mutation, but exhibited similar growth characteristics to the ancestor after heat shock. This isolate might represent a neutral mutation, although the likelihood of a neutral mutation arising, persisting, and being sampled is low. An alternate hypothesis would be that the mutation is beneficial in the context of the community it arose in due to interactions with other genotypes in the population, but not when grown in isolation.

### General tradeoff between heat shock recovery and continuous growth at elevated temperature for evolved isolates

In order to examine if the evolved isolates had any general relationship between performance in a heat shock regime (where they evolved) versus growth at an elevated temperature, we characterized the growth of all evolved isolates on methanol at 35 °C. We observed a growth defect for evolved isolates when grown at an elevated temperature of 35 °C, where almost all evolved isolates fared worse than the ancestor (Fig S4). We also observed an inverse correlation between recovery after heat shock and growth at an elevated temperature of 35 °C (Fig 5B). On average, the isolates most fit in the heat shock regime fared the worst at growth at higher temperatures, and vice versa. This effect was general and not limited to mutations in a specific locus (Table S4), suggesting a generic tradeoff between recovery after heat shock and continuous growth at an elevated temperature. In contrast, we see no such tradeoff between recovery after heat shock and growth at the optimal temperature of 30 °C (Fig S5).

### Natural isolates also display a general tradeoff between heat shock recovery and continuous growth at elevated temperature

Is the observed tradeoff between heat shock recovery and growth at elevated temperatures unique to initial adaptive steps taken in the laboratory, or is this a general pattern to be expected between strains with longer periods of selection in natural conditions? We used a panel of environmentally isolated *Methylobacterium* strains to resolve this conflict, hypothesizing that if this tradeoff is a consequence of evolution to heat shock in the laboratory, wild strains would not exhibit the tradeoff, while a more universal physiological underpinning would lead to a similar tradeoff being observed among the environmental isolates. We characterized the growth of these 9 *Methylobacterium* strains (Table S1) on methanol at either 35 °C, or recovery at 30 °C after heat shock at 55 °C for 5 min. We found the same strong tradeoff between growth after heat shock and growth at elevated temperatures (Fig 5C), suggesting competing physiological demands for growth at higher temperatures and improved heat shock resistance. We next asked whether the tradeoff is limited to methanol as a carbon source. We characterized the growth of our set of environmental isolates as before, at either 35 °C or during recovery at 30 °C after heat shock, but with succinate as the carbon source throughout. We found the tradeoff persisted with succinate as a carbon source (Fig S6). However, the carbon source used had a generic effect on the impact of heat stress, with all strains faring better under both heat shock and growth at elevated temperatures when succinate was used as the substrate instead of methanol.

## Discussion

In this work, we show *Methylobacterium extorquens* PA1 can rapidly adapt to an environmental regime of intermittent heat shock. The increases in fitness observed were driven primarily by a decrease in recovery time after heat shock, and multiple distinct genetic routes can facilitate adaptation. Despite this diversity of adaptive outcomes, we found the evolved isolates exhibited a strong tradeoff between heat shock resistance and growth at elevated temperatures. Surprisingly, this tradeoff is also present in a set of environmentally isolated *Methylobacterium* strains, indicating a general incompatibility between strategies for dealing with mildly elevated temperatures and heat shock.

A major effect of heat shock is an increased recovery time after the stress dissipates. The long lag times observed could be due to the ill-effects of heat shock and/or an active response to stressful conditions. Heat shock leads to a number of detrimental effects, such as DNA and protein damage, and ROS generation, which are known to negatively impact viability. Which of these effects drive the long recovery times is unclear. Additionally, several microbes are known to actively shut down translation in response to increased temperature via degradation of *metA*, a key enzyme in methionine biosynthesis^53^. *Methylobacterium extorquens* is also known to be capable of halting translation in response to certain stressors^54^. Further work is needed to understand the mechanisms driving long lag times following heat shock in *Methylobacterium*, and potentially other microbes. A complementary approach to understand these mechanisms would be ascertaining the molecular basis of benefits conferred by the observed mutations which decrease lag time after heat shock.

We discovered that mutations in multiple different loci are capable of improving heat shock resistance in *Methylobacterium extorquens* PA1. Beneficial mutations in multiple diverse loci are known to facilitate adaptation to growth on constant increased temperatures^19,55^. We observed a similarly large diversity of genetic targets of adaptation to heat shock. However, we do not observe a significant overlap with targets previously known to improve thermal tolerance. While a few loci have been previously known to have a role in dealing with heat shock (*dnaJ* and multiple *hsp20* loci), the links between most of the observed targets of adaptation and heat tolerance remains unclear, and warrant further study. Mutations in both *dnaJ* and *hsp20* family of proteins likely lead to increased basal expression of heat shock proteins, which is known to improve heat shock resistance^21,22^. Loss of DnaJ function aids heat shock survival in *Salmonella enterica* and *Escherichia coli* via overexpressing heat shock proteins^21^. The mutations observed in this study (Table S1) also likely lead to a loss of DnaJ activity and similarly aid heat shock survival in *Methylobacterium extorquens*. The *secA* gene encoding the protein translocase subunit was the locus with mutations in most isolates (along with DnaJ) in our evolution experiment, but its possible role in helping improve heat shock resistance remains unknown. Expression of flagellar genes has been shown to reduce stress tolerance^56^, and mutations in *flbT* (isolate F1), a regulator of flagellum synthesis^57^ might aid heat shock resistance by reducing expression of flagellum associated genes. AFG1-family ATPases have highly conserved roles across all domains of life in mediating protein degradation^58^. Heat shock is known to induce oxidative stress and protein damage, and mutations in the AFG1-family ATPase (isolate F2) likely help better deal with misfolded proteins. Mutations in cyclic nucleotide phosphodiesterase (Isolate G1) and diguanylate cyclase (Isolate C1) likely regulate the intercellular levels of key signalling molecules (e.g., cAMP, c-di-GMP) which regulate a number of core cellular processes including stress response.

Environmental variation in ecological settings is dynamic; however, most studies probing evolutionary changes in response to environmental shifts focus on constant environments. Recent studies on antibiotic resistance have found distinct evolutionary outcomes with different drug exposure dynamics^59,60^, suggesting subtle changes in the dynamics of a stressor can strongly influence evolutionary trajectories. However, it remains unclear if these differences arise due to tradeoffs between different strategies optimal for different exposure dynamics or due to differences in accessibility of different evolutionary steps in fitness landscapes which are constantly changing with different dynamics. Our results similarly suggest different dynamics of heat stress can select for distinct adaptive outcomes. Additionally, we also discovered a general tradeoff between adaptation to heat shock and growth at elevated temperatures, suggesting strategies to deal with different dynamics of heat stress can be competing. Further work is needed to understand the molecular mechanisms shaping the observed tradeoff.

Understanding the adaptive response of both individual microbes and communities to thermal stress is of great interest. A multitude of studies have greatly enhanced our understanding of the genetic basis and mechanisms which can facilitate adaptation for growth at a constant elevated temperature. However, thermal stress is not a single environment, and can manifest in distinct ways (Fig 1), each demanding distinct and perhaps incompatible strategies. Our analysis in this work focuses on two extremes among the myriad ways thermal stress can be present – constant elevated temperature and heat shock. Using evolution experiments and environmental isolates, we discover a general tradeoff between fitness in the consistently elevated temperature and heat shock regimes. While mean temperature has previously been suggested to play a role in promoting diversification of ecotypes^16,61^, our work suggests not just mean temperature, but distinct patterns of temperature fluctuations can drive or maintain diversification of *Methylobacterium* strains, both within a species and within a genus. More complex thermal gradients in nature might select for even more specialization and diversity in strategies.

Temperature is a common environmental variation faced by phyllosphere microbes like *M. extorquens*, both seasonally and daily. The temperature on the surface of leaves is not regulated and is correlated with ambient temperature, and can thus fluctuate significantly even within a day. Additionally, the relationship between leaf surface and ambient temperature is not linear, and leaf temperatures generally exceed ambient temperatures by several degrees^62,63^. This disconnect is even more prevalent for sun-exposed leaves where surface temperatures can exceed 50 °C for short periods during the day^64^. Thus, temperature variations of the kind we describe here are a common feature of life as experienced by phyllosphere microbes. An understanding of the strategies they employ to deal with, and adapt to, all aspects of temperature fluctuations is of profound importance.

Tradeoffs drive microbial diversity in ecological settings^65–67^, and this diversification fuels long-term ecological and evolutionary processes like speciation and community assembly, and shapes key cellular traits. Recent studies focusing on distinct environments have discovered key tradeoffs shaping natural microbial communities^68,69^ and fundamental cellular strategies^70–73^. In this work, we highlight how tradeoffs can also arise due to competing strategies to deal with different intensities or temporal regimes of the same stressor, high temperature. A number of recent studies successfully highlight the role of temperature in shaping microbiomes and critical ecological functions. We show that variations in temperature, and not just the direction of temperature change, can drive selection for distinct strategies. Our results suggest temperature fluctuations, even in the same direction, but of different magnitudes and durations can act as drivers of microbial diversity.

## Acknowledgements

We are very grateful to members of the Marx and Udekwu labs for discussions and helpful feedback on the manuscript. We are grateful to Tara Hudiburg for information about leaf surface temperatures, to the Martinez-Gomez lab at the University of California Berkeley for the *Methylobacterium* strains isolated from soybean and tomato leaves, and to Noah M. Arts and Alexander B. Alleman for the *Methylobacterium* strain isolated from Western Red Cedar.

## Study Funding

This work was supported by the National Science Foundation (NSF) grant DBI – 2320667 to CJM. KJR was also supported by the Hill Undergraduate Research Fellowship at the University of Idaho.

## Data Availability

Sequencing data is deposited on NCBI SRA (BioProject PRJNA1346438). All raw data and code used in this work is available on GitHub, and also in the supplementary material.

https://github.com/mallakshat/HeatShockEvolution_Tradeoff

## Notes

https://github.com/mallakshat/HeatShockEvolution_Tradeoff

### Competing Interest Statement

The authors have declared no competing interest.

