## Supplementary Information for "Evolution uncovers a general tradeoff between recovery after heat shock and growth at elevated temperatures"

for

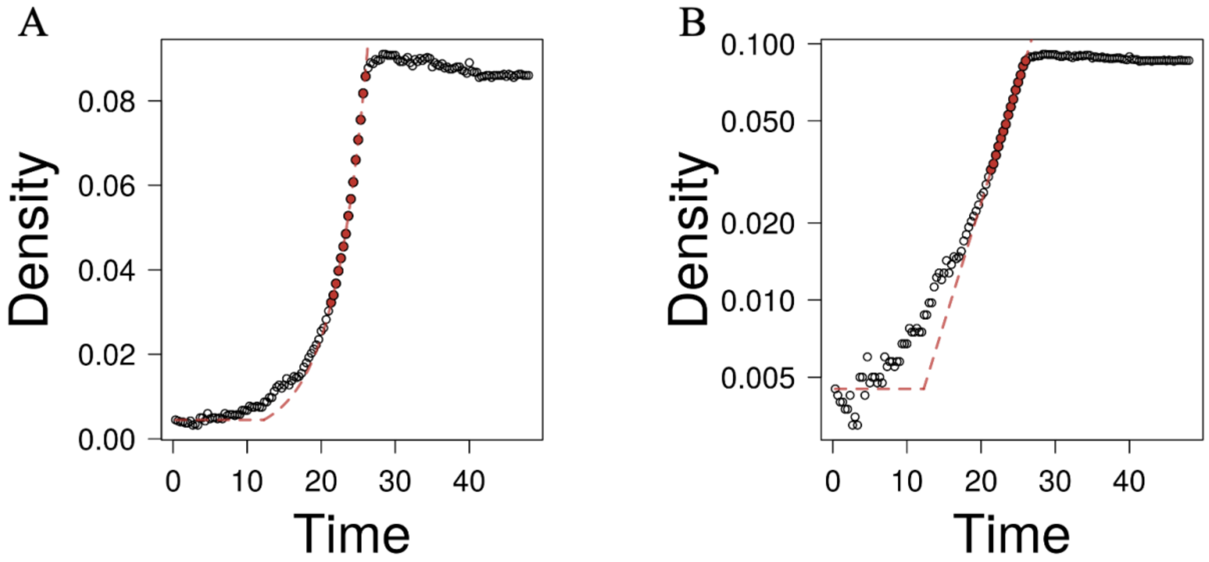


**Fig S1. Growth rates and lag times were quantified by fitting a line to log-transformed Optical Density vs time data.** Example plots of analysis shown here for one replicate of isolate F1 during recovery after heat shock. For illustration purposes, we show both **A)** regular scale, and **B)** OD axis being log-transformed. The points shaded red represent the points used to estimate maximum growth rate, and the red dashed line indicates the line of best fit for the red points, whose slope represents the maximum growth rate. The point on the time axis where this line intersects with a horizontal line from the starting OD represents the lag time. In this example, our estimated lag time = 12.26 h, and maximum growth rate = 0.216 $h^{-1}$.

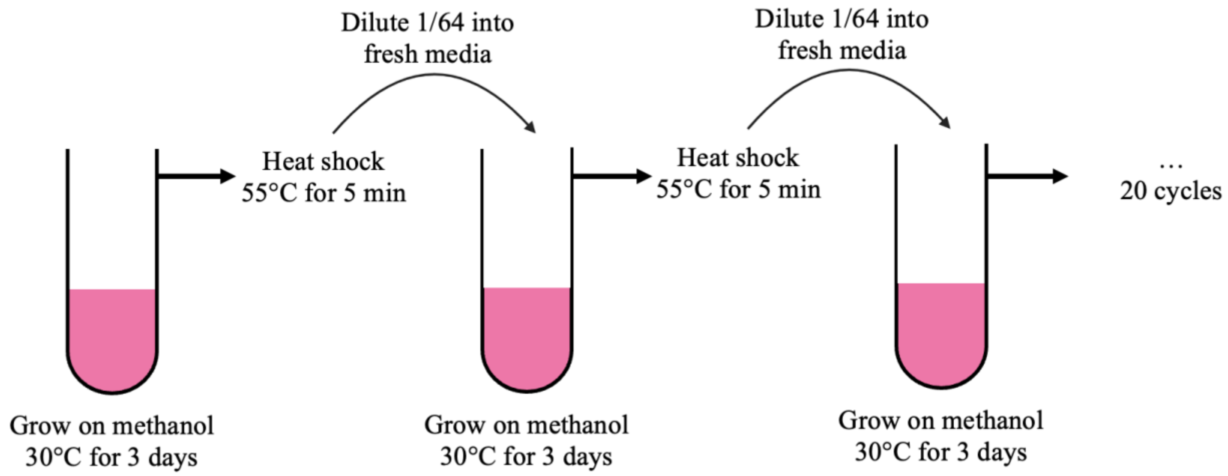


**Fig S2. Schematic of evolution experiment.** Ten replicate populations were evolved on a cyclical environment of optimal conditions (growth on methanol at 30 ˚C) and heat shock (55 ˚C for 5 min).

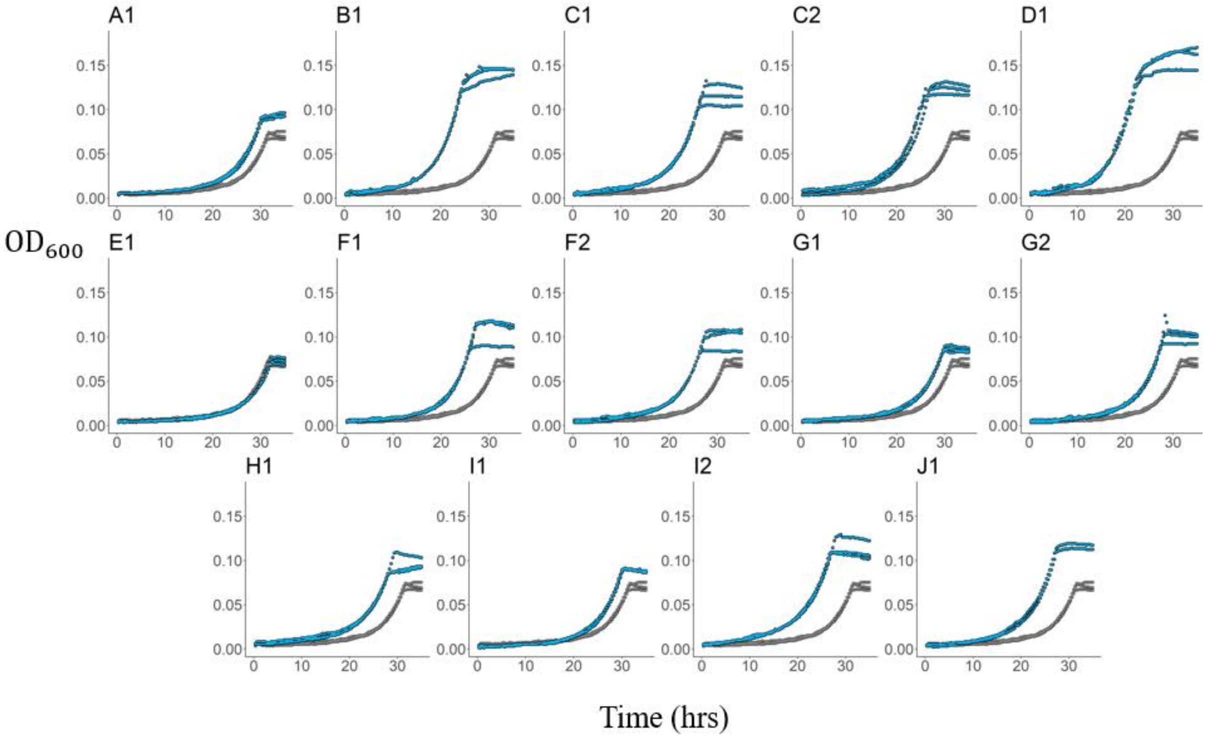


**Fig S3. Evolved isolates exhibit improved growth in the evolution regime.** Growth curves for all evolved isolates (blue) and ancestor (gray) in the evolution regime of 5 min heat shock at 55 °C followed by recovery on methanol at 30 °C. Three technical replicates shown for each isolate and the ancestor. Note that the difference in final yield is only due to substrate (methanol) evaporation with time.

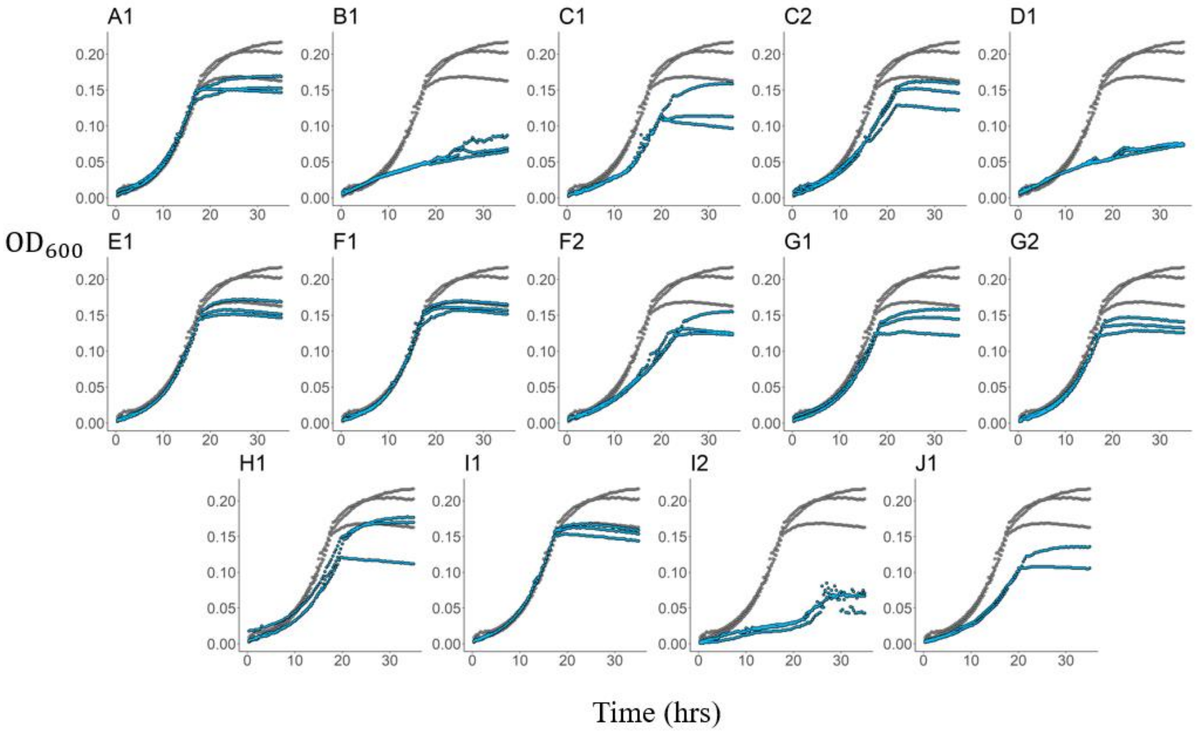


**Fig S4. Evolved isolates exhibit growth defects during growth at 35 °C.** Growth curves for all evolved isolates (blue) and ancestor (gray) during growth on methanol at 35 °C. Three technical replicates shown for each isolate and the ancestor.

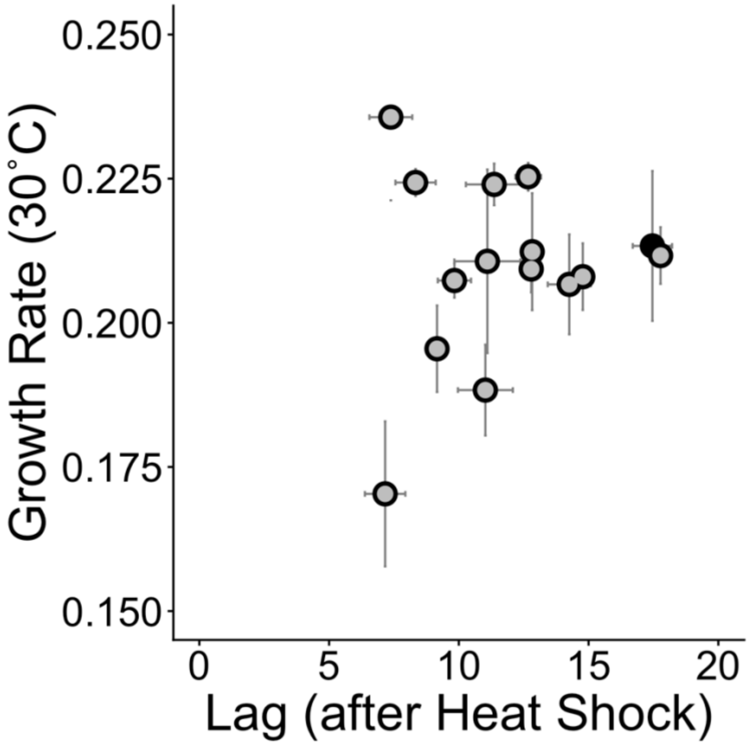


**Fig S5. Evolved isolates do not exhibit a tradeoff at the optimal growth temperature of 30˚C.** Each point represents the growth rate ($hr^{-1}$) on methanol at 30 ˚C (y-axis) vs lag time during recovery on methanol at 30 ˚C after heat shock at 55 ˚C for 5 min (x-axis) for an evolved isolate (Table 2, Table S2). The black shaded point represents the ancestor. Error bars represent standard error of mean and are visible when bigger than plotting symbols. All evolved isolates (except isolate E1) exhibit an improvement in lag time after heat shock relative to the ancestor, but they do not show any consistent change in their growth rate at the optimal temperature of 30 ˚C. This is in contrast to a strong tradeoff between improvement in a heat shock regime and fitness at a higher temperature of 35 ˚C (Fig 4B in main text).

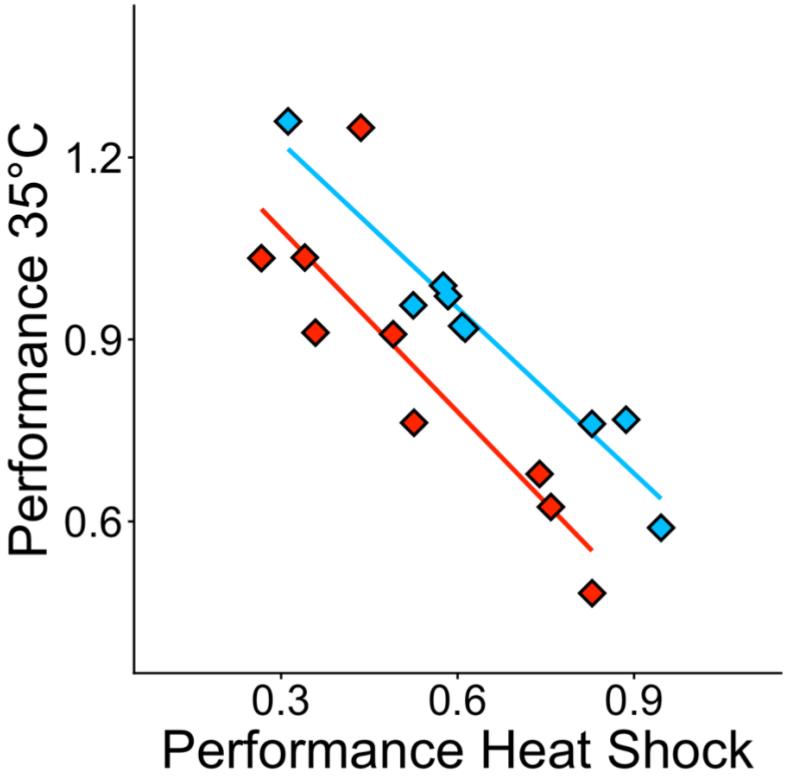


**Fig S6. Tradeoff between growth at high temperatures and recovery after heat shock for environmental isolates is independent of carbon source.** Each point represents the performance at an elevated temperature of 35 ˚C (y-axis) vs the performance during recovery at 30 ˚C after heat shock at 55 ˚C for 5 min (x-axis). We define performance as the time to reach a threshold OD in each regime relative to the time taken at an optimal temperature of 30 ˚C (Please see Fig 4A for more details on the metric). Points colored red denote results with methanol as the carbon source (same as Fig 4C). Points colored blue denote results with succinate as the carbon source. Each point represents the mean of 4 replicates. Lines represent the line of best fit for each substrate.

| Background Strain | Barcoded Strain | Environmental source |
|---|---|---|
| *M. extorquens* PA1[1] (CM2730)[2] | CM5201 | *Arabidopsis thaliana* |
| *M. nodulans* ORS 2060[3] | CM5156 | Root nodules of *Crotalaria* |
| *M. extorquens* AM1[4] (CM2720)[2] | CM5162 | Airborne contaminant |
| *M. extorquens* CM4[5] | CM5151 | Soil at a petrochemical factory |
| SLI_158* | CM5140 | Soybean leaf |
| SLI_210* | CM5141 | Soybean leaf |
| TLI_801* | CM5149 | Tomato leaf |
| TLI_802* | CM5150 | Tomato leaf |
| CM6257 [#] | n/a | Western Red Cedar leaf |

**Table S1. List of strains used to study the universality of the tradeoff (Fig 5C in main text).** All strains except CM6257 have a neutral DNA barcode inserted into the chromosome, along with a Kanamycin resistance cassette (Alexander B. Alleman, Monica J. Pedroni, Galen Beery, CJM, unpublished).

* Collen Friel and N. Cecilia Martinez-Gomez, unpublished
[#] Noah M. Arts, Alexander B. Alleman, and CJM, unpublished.

| Chromosome position | Mutation | Annotation | Gene |
|---|---|---|---|
| 1,099,002 | $(C)_{5\rightarrow4}$ | Intergenic (+63 / +28) | Mext_1011 / Mext_1012 |
| 1,289,299 | Δ1 bp | Intergenic (-281 / -257) | Mext_1169 / Mext_1170 |
| 1,520,170 | Δ7108 bp | cellulose biosynthesis deletion | Mext_1367 – [Mext_1370] |
| 1,527,312 | C → T | D239D (GAC → GAT) | Mext_1370 |
| 1,690,785 | Δ1 bp | Intergenic (-1367 / +961) | Mext_1510 / Mext_1512 |
| 3,838,949 | Δ1 bp | Intergenic (-15 / -1373) | Mext_3458 / Mext_3460 |

**Table S2. Mutations in WT ancestor (lab strain CM2730) compared to reference *M. extorquens* PA1.**

| Isolate | Annotation | Gene | Mutation | Change | Location |
|---|---|---|---|---|---|
| A1 | × | × | × | × | × |
| B1 | preprotein translocase, SecA subunit | Mext_1478 | G → A | G144D (GGC→GAC) | 1655946 |
| C1* | short-chain dehydrogenase/ reductase SDR | Mext_1019 | A → C | T220P (ACC→CCC) | 1105714 |
| | chaperone protein DnaJ | Mext_2961 | T → G | Y129D (TAC→GAC) | 3309193 |
| | diguanylate cyclase | Mext_3867 | A → T | M589K (ATG→AAG) | 4294005 |
| C2 | preprotein translocase, SecA subunit | Mext_1478 | Δ1 bp | coding (2828/2913 nt) | 1658343 |
| D1 | preprotein translocase, SecA subunit | Mext_1478 | G → T | D61Y (GAC→TAC) | 1655696 |
| E1 | N/A | Mext_3633/ Mext_3634 | A → G | intergenic | 4011581 |
| F1 | flagellar FlbT family protein | Mext_1248 | Δ1 bp | coding (345/417 nt) | 1384271 |
| F2* | AFG1–family ATPase | Mext_0934 | T → C | V239A (GTC→GCC) | 1011093 |
| | heat shock protein Hsp20 | Mext_4556 | A → G | F16L (TTC→CTC) | 5086575 |
| G1 | 3',5'-cyclic-nucleotide phosphodiesterase | Mext_1789 | Δ171 bp | coding (86-256/279 nt) | 2008936 |
| G2 | N/A (7 bp upstream of heat shock protein Hsp20) | Mext_4262/ Mext_4263 | T → A | intergenic | 4744167 |
| H1 | heat shock protein Hsp20 | Mext_1620 | G → T | K141N (AAG→AAT) | 1807080 |
| i1 | SpoVR family protein | Mext_2061 | G → C | G94A (GGC→GCC) | 2296689 |
| i2 | Chaperone protein DnaJ | Mext_2961 | Δ6 bp | coding (586-591/1158 nt) | 3309394 |
| J1* | N/A (32 bp upstream of luciferase family protein) | Mext_1738/ Mext_1739 | $(CTCG)_{6\rightarrow7}$ | intergenic | 1961229 |
| | chaperone protein DnaJ | Mext_2961 | G → A | G307D (GGT→GAT) | 3309728 |

**Table S3. Mutations identified in evolved isolates.** Isolate name denotes replicate information (A-J), and the number (1 or 2) designated to the isolate picked for that replicate. E.g. – C1 and C2 denote different individuals from the same replicate in the evolution experiment. Isolates with multiple mutations denoted with asterisks.

| Isolate | Mutation | Performance Heat Shock | Performance 35°C |
|---|---|---|---|
| A1 | × | 0.379±0 | 0.892± 0.024 |
| B1 | preprotein translocase, SecA subunit | 0.508±0 | 0.370±0.083 |
| C1* | short-chain dehydrogenase/ reductase SDR | 0.454±0.036 | 0.697± 0.01 |
| | chaperone protein DnaJ | | |
| | diguanylate cyclase | | |
| C2 | preprotein translocase, SecA subunit | 0.465±0.011 | 0.744±0.01 |
| D1 | preprotein translocase, SecA subunit | 0.572±0.058 | 0.373±0.026 |
| E1 | N/A | 0.343±0.041 | 0.839±0.025 |
| F1 | flagellar FlbT family protein | 0.44±0 | 0.868±0 |
| F2* | AFG1–family ATPase | 0.429±0 | 0.688±0.024 |
| | heat shock protein Hsp20 | | |
| G1 | 3',5'-cyclic-nucleotide phosphodiesterase | 0.381±0.025 | 0.805±0.02 |
| G2 | N/A (7 bp upstream of heat shock protein Hsp20) | 0.416±0.03 | 0.812±0.012 |
| H1 | heat shock protein Hsp20 | 0.406±0.029 | 0.745±0.036 |
| i1 | SpoVR family protein | 0.378±0.025 | 0.892±0 |
| i2 | Chaperone protein DnaJ | 0.448±0.035 | 0.418±0.007 |
| J1* | N/A (32 bp upstream of luciferase family protein) | 0.440±0.083 | 0.647±0.018 |
| | chaperone protein DnaJ | | |
| Anc | × | 0.351±0 | 0.869±0.023 |

**Table S4. Performance of evolved isolates and ancestor in a heat shock regime and during growth at 35 °C.** The tradeoff between recovery after heat shock and growth at 35 °C is not limited to mutations in a specific locus but is a general phenomenon. Values represent mean performance of 4 replicates ± standard deviation.
